# A Novel dual Influenza virus and SARS-CoV-2 neutralisation assay

**DOI:** 10.1101/2025.09.19.677384

**Authors:** Kelly A.S. da Costa, Martin Mayora-Neto, Sururat F Adigun, Titilola T Shobande, Derrick Amoaka-Tawiah, Shazeea Masud, Temitavo O Basaru-Sanni, P. Copoomootoo, Arwel Ww Jones, Glen Davison, Mariana Pjechova, Andrew D. Miller, Miroslav Fajfr, Daniel Ruzek, Nigel J. Temperton

## Abstract

Influenza and SARS-CoV-2 are co-circulating during the traditional influenza season giving rise to potential co-infections; currently estimated at 2.4%. Additionally, this means that the uptake of both vaccines (including SARS-CoV-2 boosters) will be encouraged within the same timeframe. Currently, patients receive separate vaccines in different arms on the same or different days. Induction of neutralising antibodies (NAbs) is used as the measure of efficacy for SARS-CoV-2 vaccinations while, the hemagglutination inhibition (HI) assay is the gold standard for seasonal influenza vaccine assessment. However, with the advent of novel universal influenza vaccine and dual SARS-CoV-2/Influenza vaccine approaches it is possible that functional NAb assays will become an essential component of a vaccine assessment toolbox. This assay is capable of distinguishing NAb responses during both dual vaccination development, and as part of co-infection and imprinting studies.

Taking advantage of existing pseudotyped viruses developed by us, we present herein a dual neutralisation assay, which utilises 2 different luciferase (Renilla and Firefly) reporter lentiviral vectors. We configured assays with various combinations of SARS-CoV-2 and influenza subtype/variant combinations and compared sensitivity of this dual approach to neutralisation levels seen in single virus pseudotype Micro-neutralisation (pMN) in a double-blinded test of monoclonal antibody cocktails, and subsequently with a pre-screened serum panel. Mono and dual neutralisation pMN identified correctly positive and negative samples and IC50 values were all within 2-fold of each other, suggesting that specificity and sensitivity are retained post-multiplex. As we have taken a pseudovirus based approach there is the potential to tailor to specific vaccination candidates or currently circulating strains of each virus.

The use of this versatile assay could form part of the toolbox of analysis of dual vaccination candidates and investigating responses during co-infection including studies assessing the impact of immune imprinting in the future.

## 1. Introduction

SARS-CoV-2 and Influenza are co-circulating respiratory viruses. Most countries had an influenza vaccination programme in place for the most clinically vulnerable members of society, prior to the start of the SARS-CoV-2 pandemic. Vaccine uptake for SARS-CoV-2 vaccinations has been reasonable, in fact as of August 2024 globally 70% of the global population have received at least one dose (1). Estimates of co-infection remain relatively low compared to SARS-CoV-2 mono-infection at approximately 2.4% of patients (2). However, severity of infection, damage to the lungs and mortality rate is increased during coinfection (2-5). Both influenza and SARS-CoV-2 continue to mutate and evade pre-existing immunity (6-9) and so there is continual global monitoring of the circulating variants/strain to enable the development of updated effective vaccines (10, 11). Currently, vaccinations are the best way to prevent infection and severe infections requiring hospitalisation, with the public being offered concurrent vaccination (12). This provides comparable antibody responses to those who received vaccination on separate visits (13-16). Given the lack of reported immune interference of these two vaccinations (17); to increase the number of individuals receiving both seasonal influenza and next generation SARS-CoV-2 vaccinations, one strategy currently being pursued is dual vaccination including the phase 3 Moderna trial (ClinicalTrials.gov Identifier:NCT06097273) and the Pfizer BioNtech trial (ClinicalTrials.gov identifier NTC0617899) (18-20). Innovative vaccine strategies require novel approaches to measure vaccine efficacy and tools to determine the immunological responses for a protective vaccination. The production of neutralising antibodies is used as a correlate of protection (COP) for SARS-CoV-2 (21, 22) and has been proposed as an alternative COP to HI for next generation influenza vaccines (23, 24). Live virus microneutralisation (MN) assays require specialised training, equipment and containment facilities. Pseudotype microneutralisation (pMN) assays are a viable alternative that has been shown to correlate strongly to MN (25-27) and have gained significant traction with many stakeholders during the SARS-CoV-2 pandemic. There are several advantages to using pseudotyped virus vs. the wildtype; they are a safer, with lower containment requirements, flexible and relatively rapid method to test antibody responses against a range of influenza subtypes/SARS-CoV-2 variants which have already been developed by ourselves and others (28-33) and have multiple uses (4, 28, 34, 35). If evaluating neutralising antibody responses to the two viruses in a dual vaccination setting, it makes sense that an assay, which can measure these within the same sample, would be a valuable tool for vaccine immunogenicity assessment. Herein, we demonstrate for the first time that it is possible to detect neutralising antibody responses (NAbs) to influenza and SARS-CoV-2 in the same sample using our existing lentivirus pseudotyping system for influenza and SARS-CoV-2 which builds on a previous report showing we can detect two separate subtypes of influenza (36, 37).

## 2. Materials and Methods

### 2.1 Cell Culture

For production of SARS-CoV-2 and Influenza pseudotyped viruses (PV), human embryonic kidney 293T/17 (HEK293T/17, ATCC: CRL-11268^a^) were maintained in Dulbecco’s Modified Essential Medium (DMEM); DMEM (PANBiotech P04-04510) supplemented with 10% (v/v) heat-inactivated Fetal Bovine Serum (PANBiotech P30-8500), and 1% (v/v) Penicillin-Streptomycin (PenStrep) (Sigma P4333)) at 37°C and 5% CO_2_.

Target cells for assay were prepared by transiently pre-transfecting HEK293T/17 cells with 2000ng Angiotensin Converting Enzyme 2 (ACE2) plasmid (pCDNA3.1+) (38) and 150ng of Transmembrane Serine Protease 2 (TMPRSS2) plasmid (pCAGGS) (39) using Fugene HD. HEK293T/17 cells were passaged in a T75 flask 1:8 the day before transfection and then used 24 hrs after transfection.

### 2.2 Plasmid production and transformation

Influenza hemagglutinin (HA) and neuraminidase (NA) genes from influenza A (IAV) H1N1 (A/England/195/2009) & H3N2 (A/SouthAustralia/34/2019) and influenza B (IBV) B/Victoria-like (B/Washington/2/2019) were cloned into pEVAC plasmids (GeneArt). All HA and NA genes were codon-optimised and adapted to human codon use using the GeneOptimiser algorithm (32, 40, 41). SARS-CoV-2 (Wuhan) spike expression plasmid (pCAGGS) was obtained from CFAR, (catalogue no. 100976), SARS-CoV-2 Delta (B.1.617.2) variant was a gift from Gavin Screaton, University of Oxford, Omicron (BA.1) was a gift from G2P, both variants encoded in pcDNA3.1 expression vector.

Plasmids were transformed in competent *E*.*Coli* DH5α cells (Invitrogen 18265-017) using the heat-shock method as preciously described (32, 41). pCAGGS and pcDNA3.1 plasmids confer ampicillin (AMP) resistance and pEVAC confers Kanamycin resistance. Plasmid DNA was extracted using QIAprep Spin Miniprep Kit (Qiagen 27104) and quantified using UV spectrophotometry (NanoDrop™ Thermo Scientific) and stored at -20°C for later use.

### 2.3 Pseudotyped Virus Production

Influenza HA NA Pseudotyped Viruses (PV) were produced via four-plasmid co-transfection as described previously (42). Briefly, 4×10^5^ HEK 293T/17 cells in complete DMEM were seeded per well of a 6-well plate and incubated at 37°C, 5% CO_2_ overnight. The next day, media was replaced and cells were transfected using Opti-MEM™ (Thermo Fisher Scientific 31985062) and FuGENE® HD Transfection Reagent (ProMega E2312) with the following plasmids: 10ng HA encoding plasmid (pEVAC), 5ng NA encoding plasmid (pEVAC), 375ng luciferase reporter plasmid Firefly or Renilla (43), and 250ng p8.91 gag-pol (Gag-Pol expression plasmid (44, 45)). A protease plasmid was included: for H1N1 (A/England/195/2009), 5ng type II Transmembrane Protease Serine 4 (TMPRSS4), and for H3N2 (A/Switzerland/8060/2017) and BVIC (B/Washington/2/2019), 5ng human airway trypsin-like protease (HAT) as previously determined (32). SARS-CoV-2 PVs were produced using 3 plasmid co-transfection as previously described (31) and represented graphically (supplementary figure 1 (s1)) Briefly T75 cell culture flasks were seeded at 2×10^5^ cells/mL and incubated at 37°C, 5% CO_2_ overnight. The next day, media was replaced and cells were transfected using Opti-MEM™ (Thermo Fisher Scientific 31985062) and FuGENE® HD Transfection Reagent (ProMega E2312) with the following plasmids: 1µg Spike encoding plasmid, 1µg p8.91 gag-pol (Gag-Pol expression plasmid (44, 45)) and 1.5µg of luciferase reporter plasmid (Renilla or firefly). All plasmid DNA were combined in OptiMEM and incubated with FuGENE® HD for 15 minutes. The plasmid DNA-OptiMEM mixture was then added to the cells with constant swirling. Forty-eight hours post-transfection, supernatant was collected, passed through a 0.45 μm filter and stored at −80°C. Plates were incubated at 37°C, 5% CO_2._

### 2.4 Test panel and serum samples

A blinded test panel was created to test if the dual and mono pMN assays can distinguish positive and negative controls correctly as described in table 1 below and serially diluted 1:2 from an initial dilution of 1:100. Reference antisera to Influenza A and B were obtained from the National Institute for Biological Standards and Control (MHRA-NIBSC). The SARS-CoV-2 monoclonal antibody used was a gift from Nanotools (46). Archived sera samples collected in 2011/2012 as part of a previous study (47) were used as pre-pandemic (SARS-CoV-2 negative) samples. Sera samples collected from 5 healthy SARS-CoV-2 vaccinated controls and 5 individuals infected during the first wave of SARS-CoV-2 in the Czech Republic as per local ethics regulations. The 5 samples in each group were selected at random. All Sera was obtained by centrifuging serum separator tubes at 400g for 5 mins and stored at -20°C in the UK or shipped to the UK on dry ice and stored at -20°C for further analysis.

**Table 1.** Positive and negative control panel. A consisting of a universal negative control (FBS), Influenza positive controls for each subtype being tested (A/H1N1, A/H3N2 & B/Victoria) in the form of anti-sera purchased from MHRA (NIBSC, UK), a universal positive control was made from Influenza and SARS-CoV-2 positive controls and was also influenza subtype specific.

|  |  |  |
| --- | --- | --- |
| 1 | Negative control | FBS |
| 2 | Influenza positive control (subtype specific anti-sera) | a) H1 – anti-A/Puerto Rico/8/1934<br>b) H3 – anti- A/Darwin/9/2021<br>c) BVIC – anti-B/Brisbane/60/2008 |
| 3 | SARS-CoV-2 positive control | SARS-CoV-2 Monoclonal antibody from Nanotools (46) |
| 4 | Positive for Influenza and SARS-CoV-2 | 50% Influenza positive control & 50% SARS-CoV-2 positive control |

### 2.5 Titration of Pseudotyped Viruses

The production titre of the PVs used was determined as previously described (31, 32). Briefly, PV transduction titres were determined by transduction of target cells (prepared as described 2.1). 50µl of 2-fold serially diluted Influenza and SARS-CoV-2 PVs were prepared in flat-bottomed white 96 well plates (Thermo Scientific 10072151). Target cells were added to PVs at 3.5x 10^5^ cells/mL, including to cell only wells. Plates were incubated for 48h at 37°C, 5% CO_2_ after which all media was removed from all wells and cells were lysed with Bright-Glo® reagent or Renillia-Glo® reagent (Promega). Luminescence was measured using a GloMax Navigator® (Promega) with the Glomax® luminescence Quick Read protocol or GloMax® Renilla Luminescence quick read programme. Titres were calculated as RLU/mL using Microsoft Excel.

### 2.6 Mono and dual pseudotyped virus micro-neutralisation assays (pMN)

pMN assays were performed as previously described (31, 48). Briefly, the blinded panel samples or the human sera were diluted 2-fold from 1:40 initial dilution in a white 96 well plate (Thermo Fisher Scientific 136101). For mono pMN assays a single virus was diluted to achieve approximately 1×10^6^ RLU/well and for dual pMN two viruses were diluted to achieve approximately 1×10^6^ RLU/well for each virus. In both assays an equal volume of PV was added to each well and incubated for 1 hour at 37°C and 5% CO_2_, before the addition of target cells (HEK293T/17 cells transiently transfected to express TMPRSS2 and ACE2) at 3 × 10^5^ cells/mL. Plates were incubated for 48h at 37°C, 5% CO_2_ after which all media was removed from all wells, and cells were lysed with either Bright-Glo® reagent, Renillia-Glo® reagent (Promega) or Dual-Glo® reagent as required. Luminescence was measured using a GloMax Navigator® (Promega) with the Glomax® luminescence Quick Read protocol, the GloMax® Renilla Luminence quick read programme or the Dual Glo® luminescence protocol. Finally, IC_50_ values were determined by non-linear regression using GraphPad Prism (v.10) as previously described (49). All experiments were performed on 3 separate occasions with samples in duplicates.

### 2.5 Statistical analysis

All statistical analyses were performed with GraphPad Prism 10 for Windows (GraphPad Software). pMN titres were normalised and IC_50_ values were calculated by dose response inhibition analysis. Mann-Whitney (non-parametric) testing was used to compare data and Spearman’s non-parametric correlation analysis was used for comparison of Mono and dual neutralisation assays.

## 3. Results

### 3.1 Pseudotyped virus production with Renilla and Firefly luciferase

A representative strain from Influenza A and B subtypes were used to reflect those in the trivalent influenza vaccine were selected i.e. H1N1, H3N2, & Influenza B Victoria-like linage (B/Washington/2/2019). SARS-CoV-2 PVs were produced to express the Spike protein from Wuhan, Delta (B.1.617.2) and Omicron (BA.1) variants. Both influenza and SARS-CoV-2 PVs were produced with a Renilla luciferase reporter and a firefly luciferase reporter (figure S1). Production titres (RLU/mL) of Influenza and SARS-CoV-2 PVs produced with Renilla reporter were not significantly different from titres using firefly luciferase (figure S2). However, the mean production titre of IAV PV H3N2 was higher when produced with firefly luciferase (figure S2) and so the final assay was developed with firefly luciferase reporter for influenza strains and Renilla luciferase was used as the reporter for the SARS-CoV-2 PVs.

### 3.2 Panel test – mono and dual pseudotyped neutralisation assays

To confirm that the assay was capable of detecting virus specific neutralisation, a blinded panel was prepared as described in table 1. This panel contained one universal negative sample, one influenza neutralising anti-sera (subtype specific) and one SARS-CoV-2 MAb with differing levels of neutralising activity against the variants used herein. The panel was tested with both the traditional mono pMN and compared to the dual pMN using the same luciferase reporters for each PV. The negative remained negative (IC50 =0) when using both the dual and mono-neutralisation assays. Similarly, single neutralising antibodies only neutralised the specific viral target regardless of the presence of a second PV in the well (figure 1). The IC50 values obtained in dual pMN were slightly lower than those obtained with the mono pMN but all within 2-fold of the mono pMN (Figure 1C) and these IC50 values correlate (Spearman correlation p= 0.0001) (figure 2D).

**Figure 1.**
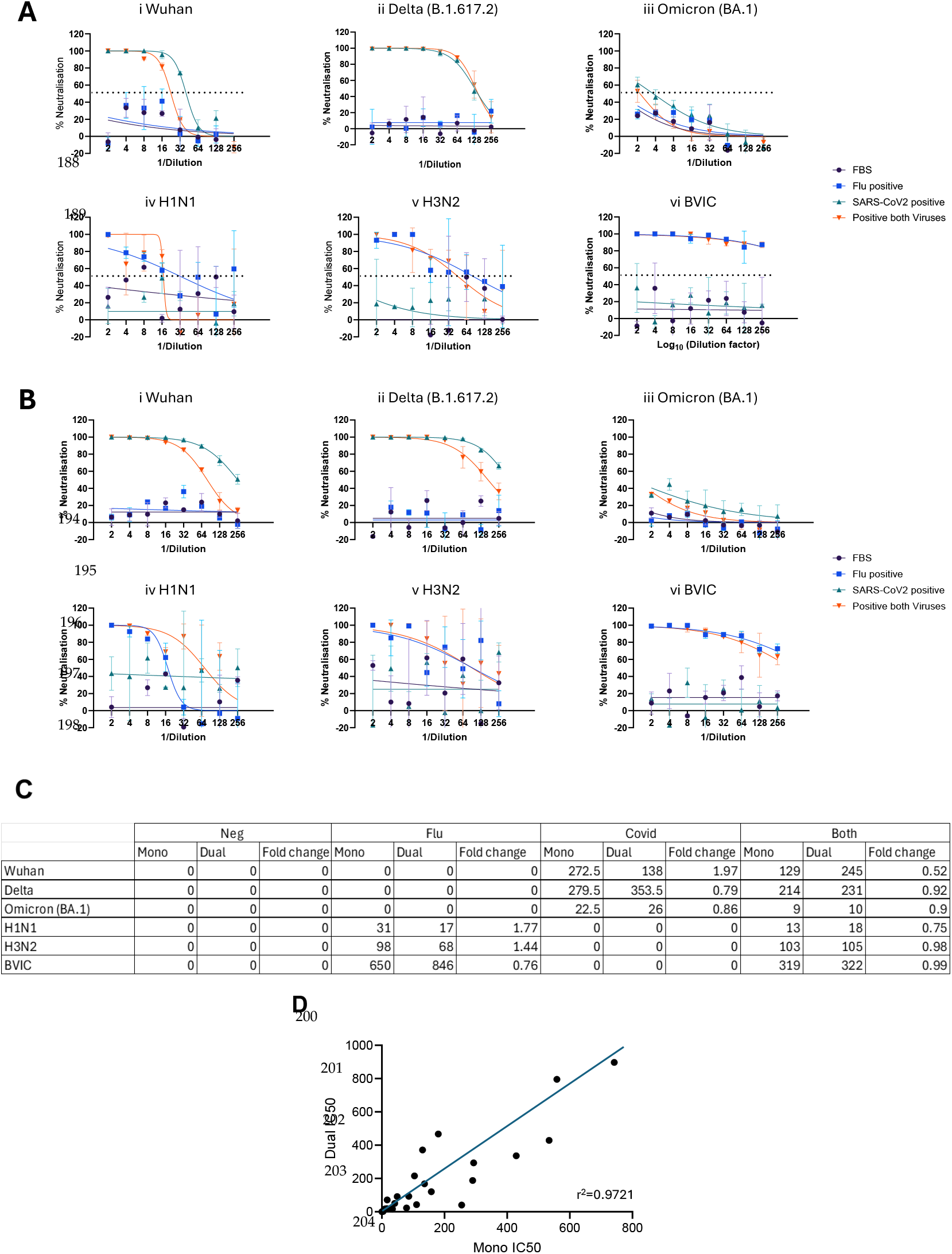
Comparison of mono and dual pMN assay with test panel. Mono and dual pMN were performed with a blinded panel (table 1). Experiments were carried out in duplicate on 2 separate occasions and the mean percentage neutralisation and IC50 were calculated (± SD). **A** mono pMN with SARS-CoV-2 variants (i) Wuhan, (ii) Delta and (iii) Omicron (BA.1) and Influenza subtypes (iv) H1N1 (A/England/195/2009), H3N2 (A/SouthAustralia/34/2019), BVic (B/Washington/2/2019) and **B** dual pMN (i) Wuhan, (ii) Delta and (iii) Omicron (BA.1) (iv) H1N1 (A/England/195/2009), H3N2 (A/SouthAustralia/34/2019), BVic (B/Washington/2/2019). **C** The evaluation of dual pMN was carried out by comparison of IC50 and calculation of fold-change from results obtained with mono pMN assays (2d.p). **D**. Comparison of IC50 titres obtained using dual and mono-neutralisation. Scatterplot showing the correlation of IC50 values from the same samples using mono pMN and Dual pMN. Spearman correlation gave a P value <0.0001.

**Figure 2.**
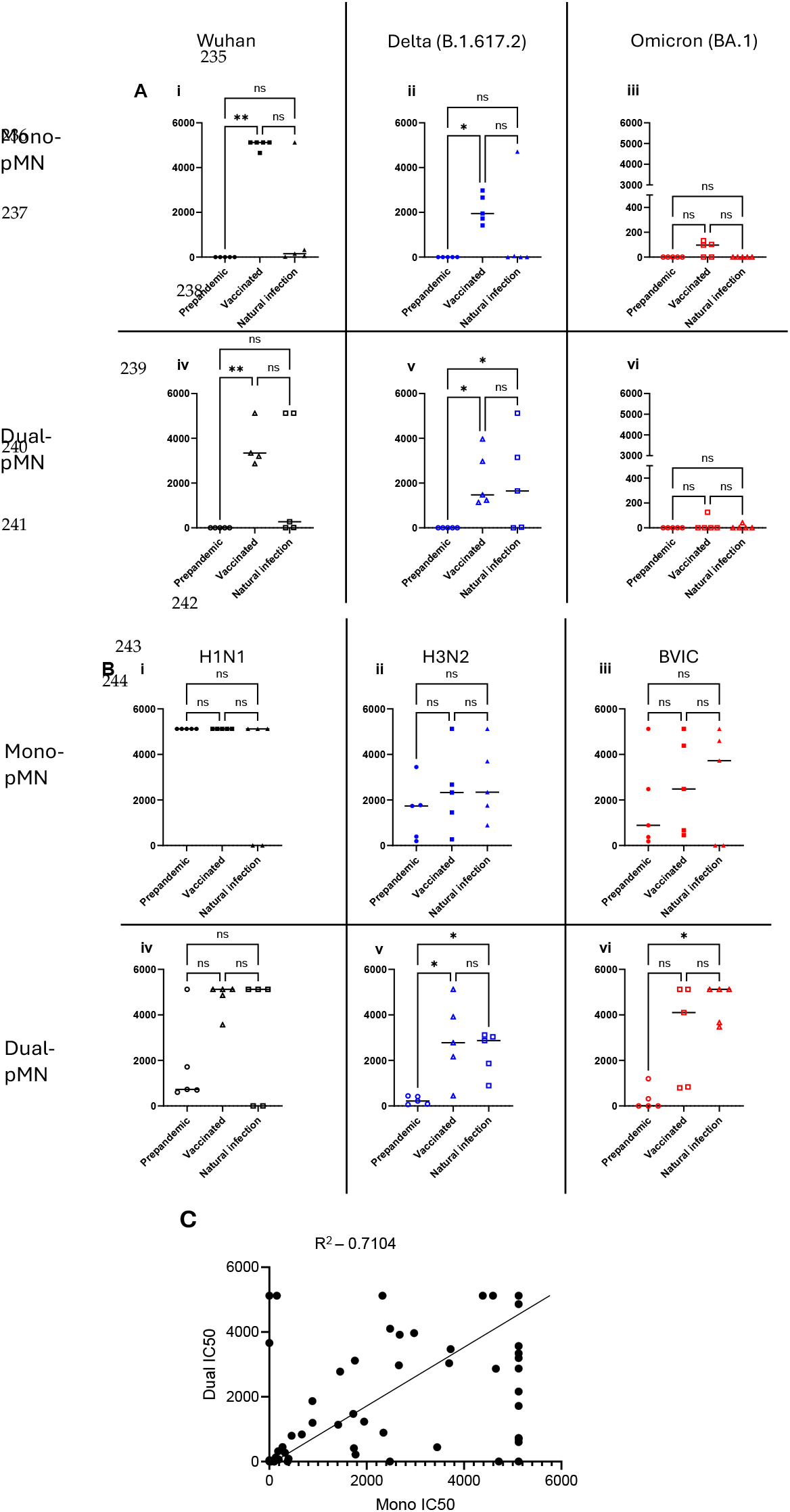
Comparison of mono and dual pMN assay with human sera test panels. Mono and dual pMN were performed with 3 groups of human sera 1 – pre-pandemic sera, 2 – Healthy volunteers vaccinated with SARS-CoV-2 in Czech Republic and 3 – individuals infected with SARS-CoV-2 during the first wave in the Czech Republic (section2.4). Experiments were carried out in duplicate on 2 separate occasions and the mean percentage neutralisation and IC50 were calculated (± SD). **A** mono-pMN with SARS-CoV-2 variants (i) Wuhan, (ii) Delta (B.1.617.2) and (iii) Omicron (BA.1) and dual-pMN (iv) Wuhan, (v) Delta (B.1.617.2) and (vi) Omicron (BA.1) and **B** mono-pMN with Influenza subtypes (i) H1N1 (A/England/195/2009), (ii)H3N2 (A/SouthAustralia/34/2019) and (iii) BVic (B/Washington/2/2019) and dual pMN (iv) Wuhan, (v) Delta (B.1.617.2) and (vi) Omicron (BA.1). **C** Comparison of IC50 titres obtained using dual and mono-neutralisation. Scatterplot showing the correlation of IC50 values from the same samples using mono pMN and Dual pMN. Spearman correlation gave a P value <0.0001.

### 3.3 Human serum test – mono and dual neutralisation assays

We subsequently tested 3 groups of blinded sera samples, one set of pre-pandemic samples and two sets of post-pandemic samples. 5 samples from vaccinated individuals and 5 samples from individuals infected with SARS-CoV-2 during the first wave of the pandemic in the Czech Republic before they received the SARS-CoV-2 vaccination. We used these panels to determine the levels of antibodies to various strains of influenza /variants of SARS-CoV-2 present from vaccination or infection. Furthermore, these sera were used to compare neutralising antibody titres (IC50) obtained when using dual-pMN and mono-pMN. All sera were tested using both assays on the same day and repeated on 2 separate occasions. The pre-pandemic samples were all found to be negative for all participants for all the SARS-CoV-2 variants tested, and this was the same when either mono or dual neutralisation pMN assays were used (figure 2A). Following vaccination with the first generation of SARS-CoV-2 vaccines, or natural infection during the first wave of the pandemic, the highest levels of neutralising antibody titres (IC50) were obtained against Wuhan strain (figure 2A); again, this trend was seen when using both mono-pMN or dual-pMN. Significant differences were seen in antibody IC50 titres to Wuhan and Delta (B.1.61.2) variants in post SARS-CoV-2 vaccination or natural infection groups compared to pre-pandemic group sera (Figure 2A i,ii,iv & v). Whereas, as expected, the IC50 titres to the later SARS-CoV-2 variant Omicron (BA.1) were low or undetectable regardless of whether the dual or mono-pMN assay was used (Figure 2A iii &vi). The neutralising antibody titres against Influenza H1N1 were not significantly different between the sera groups (Figure 2B i & iv). Interestingly there was a trend towards an increase in the NAb titre after the pandemic against Influenza A H3N2 and B Victoria (Figure 2B ii, iii, v, iv) however, the influenza vaccination and infection history is unknown for these individuals. There were small differences between the dual- and mono-pMN assays but all were within 2-fold difference of each other (figure 2) and these IC50 values correlate (Spearman correlation p= 0.001) (figure 3C).

## 4. Discussion

Current advice for clinically vulnerable people and front-line workers is to be vaccinated against influenza and SARS-CoV-2 on an annual basis (50). Concurrent vaccination in the same or different arms have been explored and studies have indicated that comparable levels of neutralising antibodies are obtained with both approaches (12-16)

Neutralising antibody responses have been shown to correlate well to the effectiveness of currently SARS-CoV-2 licensed vaccines (51, 52). However, the gold standard correlate of protection (COP) for influenza vaccine efficacy is hemagglutination inhibition assay (HI) titre. There is scope to look for NAb responses against influenza during vaccine development particularly with universal vaccine approaches which target sites other than the head of the HA protein (52, 53). Momentum for the development of a universal influenza vaccine which can protect against multiple subtypes of influenza has increased following the transmission of influenza A H5 in dairy cows in the USA (54, 55) and several other species of mammals (globally) in south America (56-58). Additionally, various broadly neutralising antibodies (bNAbs) have been identified which target the stem region of HA (HA2) of influenza and several bNAbs target spike of SARS-CoV-2 (and potentially other coronaviruses) which could be used as treatments or as prophylaxis (46, 59, 60). Therefore, we propose that at least in developmental stages it is useful to use NAbs as an indicator of which vaccine candidates would be worth pursuing in the search for a dual vaccination against both SARS-CoV-2 and Influenza. The pMN assay has been used extensively by us and others in SARS-CoV-2 research since 2020 (61-64). As with Influenza pseudotyped viruses this means that strains/variants can be investigated at lower levels of containment, and that as new strains/variants are identified PVs can be updated relatively quickly via SDM or de novo synthesis (32, 61-65).

Herein, we have shown that it is possible to detect neutralising antibody responses to two viruses from two different families in one well, Influenza A or B and a SARS-CoV-2 variant. This assay has been constructed using PVs with two distinct luciferases as reporters, Firefly and Renilla, and takes advantage of commercially available reagents. This work builds on a previous study by us, which showed that this system can be tailored to detect two subtypes of HPAI Influenza A (H5 and H7) (36, 37). We further modified the established pMN by increasing the concentration of cells to ensure that a sufficient number of targets were available.

Using a blinded panel we have shown that it is possible to detect the presence of neutralising antibodies to both or either virus. Moreover, we have shown that IC50 titres obtained using the mono and dual pMN assays are within 2-fold of each other and these titres correlate strongly with each other. Using the dual pMN we also demonstrated comparable results with a panel of sera from individuals collected pre- and post-the SARS-CoV-2 pandemic and stratified by SARS-CoV-2 vaccination and infection status. Data obtained here indicates that NAb titres of patients vaccinated with first generation vaccine or infected during the first wave of SARS-CoV-2 is lower when tested against later VOCs such as Omicron (BA.1 here). This concurs with published data (66-69). The group tested here was small and the pre- and post-pandemic samples have not been age matched. However, there is a trend in the samples tested to have a higher H3 and Bvic responses in the post-pandemic group. There is some evidence that immunodynamics to influenza have been altered following the SARS-CoV-2 pandemic (70, 71) and this is an avenue of research which would be interesting to pursue outside the scope of this study, but given the limited availability of pre-pandemic samples the dual neutralisation assay could be useful. Overall, the dual pMN assay presented here is flexible, accurate and can be used to distinguish neutralising antibody responses to Influenza and SARS-CoV-2 from a single serum sample. This means that various combinations of responses can be configured, and these can be updated depending on the vaccine and circulating strains/variants. There is scope to adapt this assay to different viruses, and this could be applied to other sero-surveillance for example in animals where sampling is more difficult as minimal amounts of sera are obtained in addition to dual vaccine candidates. We have demonstrated the application of this assay to contribute to vaccine design, sero-surveillance and to determine strain specific infections to inform clinical treatments offered to co-infected patients.

## Author Contributions

Conceived and designed experiments – KdC & NT; Performed experiments – KdC, SFA DAT SM TOBS JC TTS; Plasmid construction: MMN KdC; Analysed the data – KdC, SFA; Reagent and sample provision – NT JH ADM MF DR, AWJ GD; Wrote the paper – KdC; Revised and edited the paper – KdC, MMN, ADM MF GD AWJ MP DR & NT

## All authors reviewed and approved the final manuscript

## Acknowledgments

We would like to thank all participants who donated sera, G2P and Nanotools for the provided reagents.

