## Supplemental Figures for "A Novel dual Influenza virus and SARS-CoV-2 neutralisation assay"

### Supplementary Material

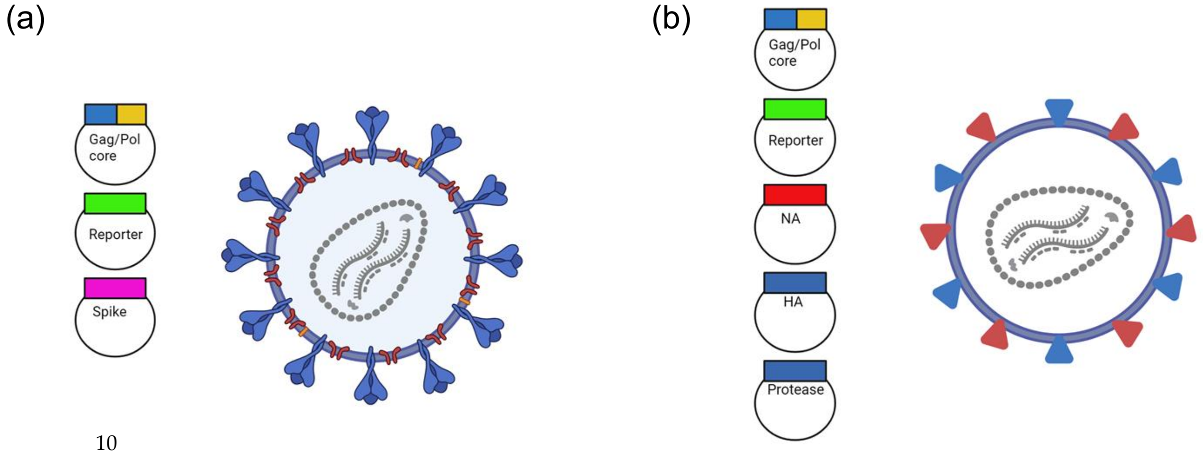


**Supplementary Figure 1. (S1) Construction of lentiviral pseudotyped viruses (PVs) for use in dual neutralisation assay.** (a) SARS-CoV-2 spike (S) expressing PV constructed with a Renilla luciferase reporter plasmid. PVs were made expressing S from Wuhan, Delta (B.1.617.2) and Omicron (BA.1) variants and (b) Influenza hemagglutinin (HA) and neuraminidase (NA) expressing PVs constructed with a firefly luciferase reporter plasmid. Influenza PVs were constructed with strain matched Influenza A H1N1, H3N2 & Influenza B Victoria-like linage.

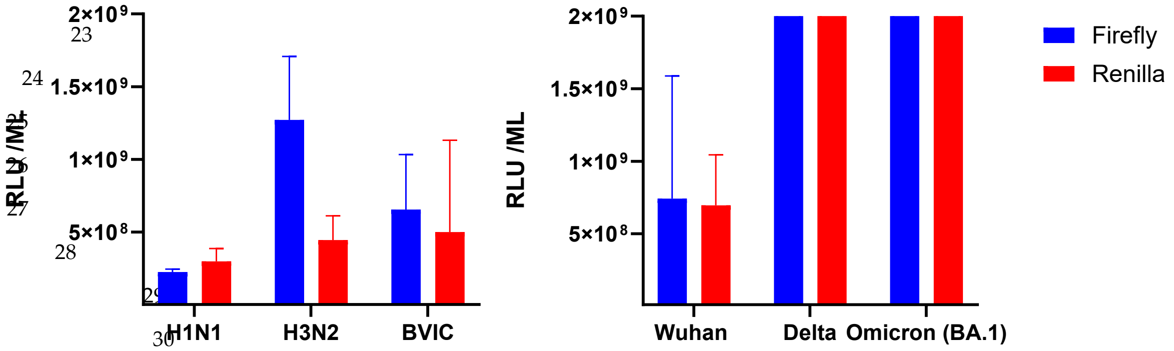


**Supplementary Figure 2. (S2) Comparison of mean titres** obtained with Renilla and Firefly luciferase for Influenza and SARS-CoV-2 PVs (n=4 batches). No significant difference observed between titres obtained with either luciferase.

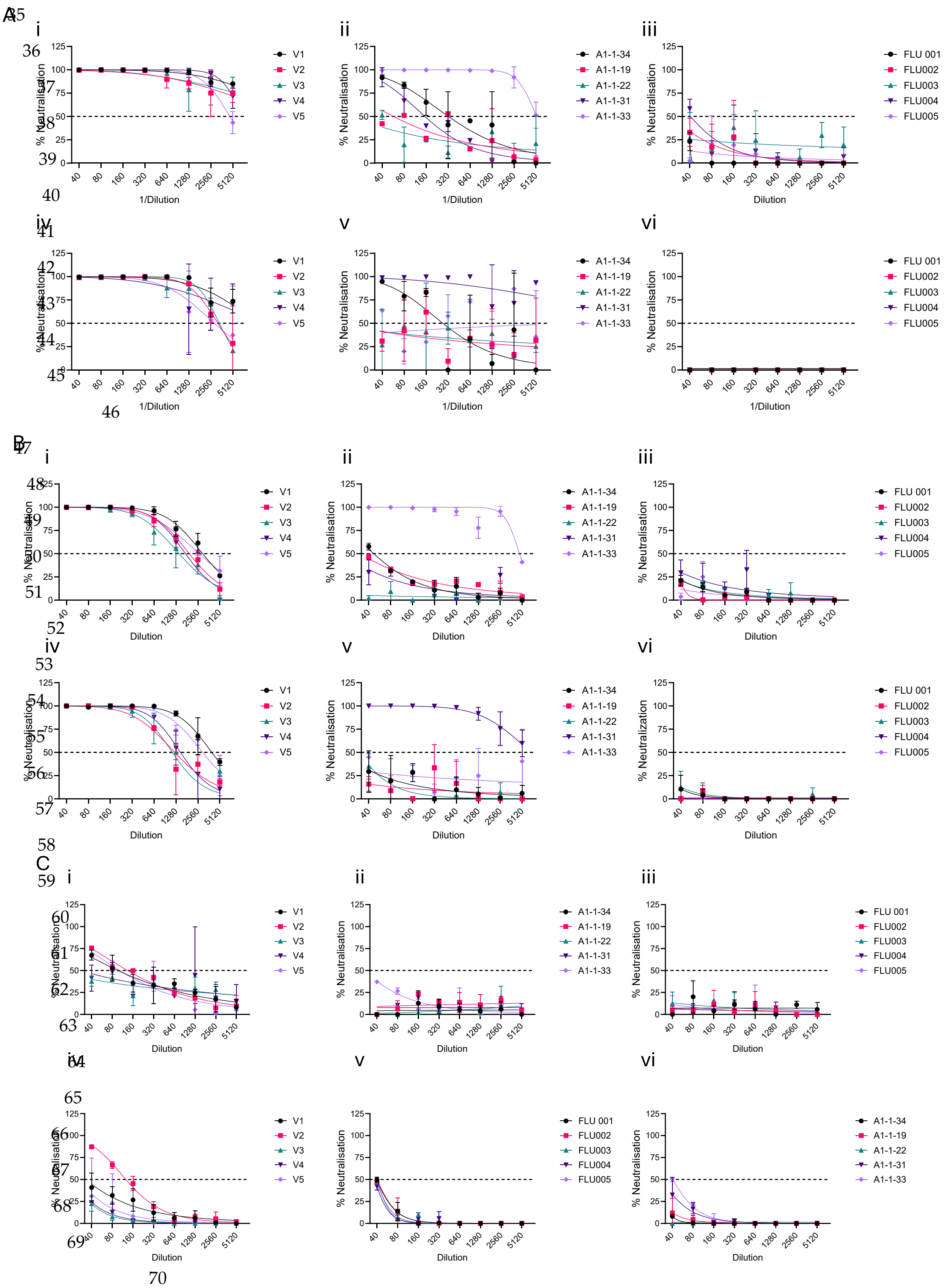


**Supplementary Figure 3. (S3) 3 groups of 5 patient sera were tested with mono-pMN SARS-CoV-2 and dual pMN SARS-CoV-2/Influenza assays.** All samples were tested on 2 separate occasions. Patients sampled before the pandemic (pre-pandemic group) (Flu001 – Flu 005), following vaccination with 1st generation SARS-CoV-2 vaccines (V1-V5) and following natural infection during the first wave of SARS-CoV-2 in the Czech republic. A- SARS-CoV-2 Wuhan variant (i-iii) mono-pMN and dual-pMN with H1N1 influenza. B- A- SARS-CoV-2 Delta (B.1.617.2) variant (i-iii) mono-pMN and (iv-vi) dual-pMN with A/H3N2 influenza, C- SARS-CoV-2 Omicron (BA.1) variant (i-iii) mono-pMN and (iv-vi) dual-pMN with B/Victoria influenza. All samples were tested on 2 separate occasions in duplicate (±SD).

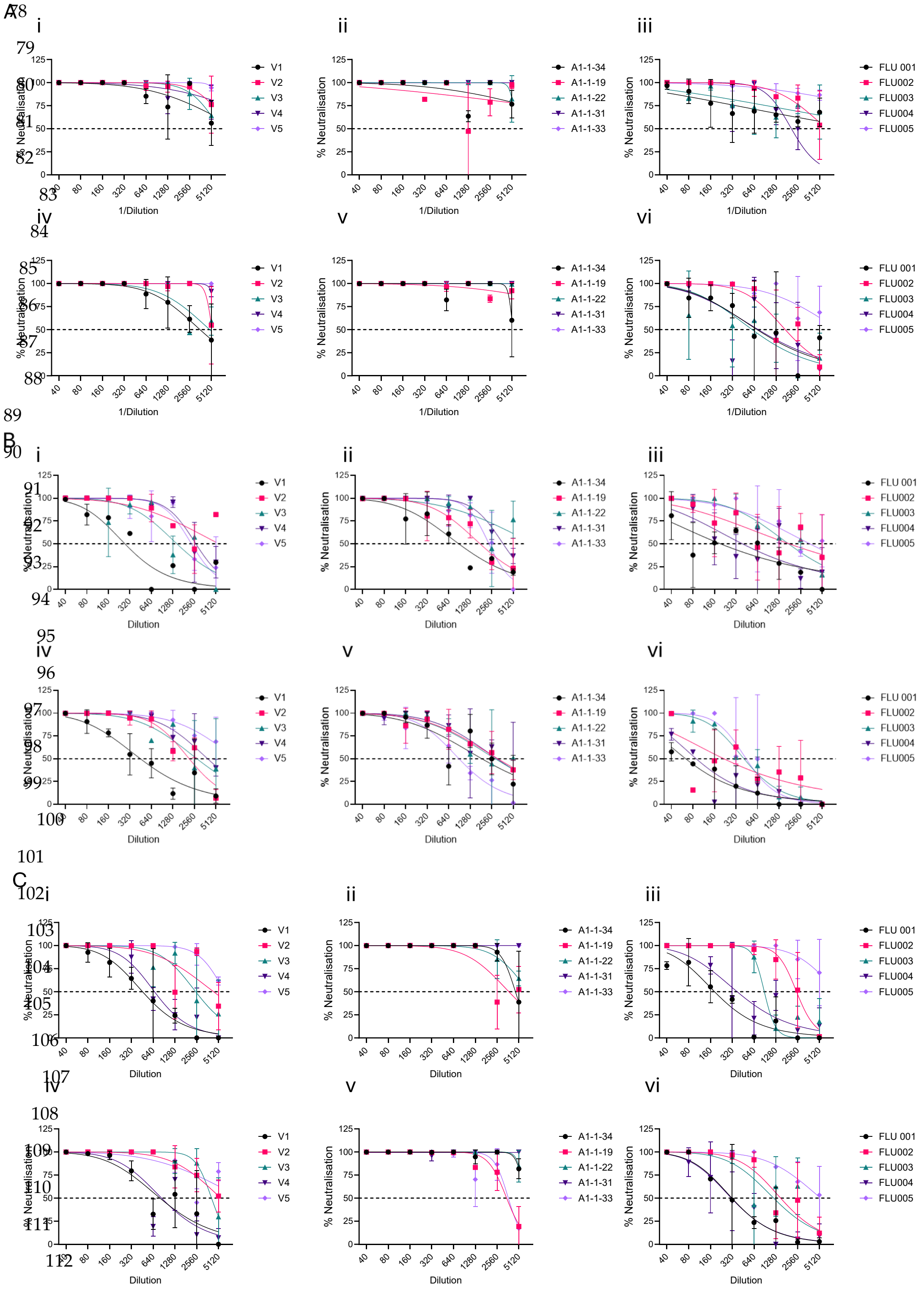


**Supplementary Figure 3. (S3) 3 groups of 5 patient sera were tested with mono-pMN Influenza and dual pMN SARS-CoV-2/Influenza assays**. Patients sampled before the pandemic (pre-pandemic group) (Flu001 – Flu 005), following vaccination with $1^{st}$ generation SARS-CoV-2 vaccines (V1-V5) and following natural infection during the first wave of SARS-CoV-2 in the Czech republic. A- Influenza A H1N1 (i-iii) mono-pMN and dual-pMN with SARS-CoV Wuhan variant B- Influenza A H3N2 (i-iii) mono-pMN and (iv-vi) dual-pMN with SARS-CoV-2 Delta (B.1.617.2) variant, C- Influenza B Victoria lineage (i-iii) mono-pMN and (iv-vi) dual-pMN with SARS-CoV-2 Omicron (BA.1) variant. . All samples were tested on 2 separate occasions in duplicate (±SD).
